# Comparative genomics among three cyst nematode species reveals distinct evolutionary histories among effector families and an irregular distribution of effector-associated promoter motifs

**DOI:** 10.1101/2021.11.01.466756

**Authors:** Joris J.M. van Steenbrugge, Sven van den Elsen, Martijn Holterman, Jose L. Lozano-Torres, Vera Putker, Peter Thorpe, Aska Goverse, Mark G. Sterken, Geert Smant, Johannes Helder

## Abstract

Potato cyst nematodes (PCNs), an umbrella term used for two species, *Globodera pallida* and *G. rostochiensis*, belong worldwide to the most harmful pathogens of potato. Pathotype-specific host plant resistances are an essential handle for PCN control. However, the poor delineation of *G. pallida* pathotypes hampers the efficient use of available host plant resistances. Long-read sequencing technology allowed us to generate a new reference genome of *G. pallida* population D383 and, as compared to the current reference, the new genome assembly is 42 times less fragmented. For comparison of diversification patterns of six effector families between *G. pallida* and *G. rostochiensis*, an additional reference genome was generated for an outgroup, the beet cyst nematode *Heterodera schachtii* (IRS population). Large evolutionary contrasts in effector family topologies were observed. While VAPs diversified before the split between the three cyst nematode species, the families GLAND5 and GLAND13 only expanded in PCN after their separation from the genus *Heterodera*. Although DNA motifs in the promoter regions thought to be involved in the orchestration of effector expression (‘DOG boxes’) were present in all three cyst nematode species, their presence is not a necessity for dorsal gland-produced effectors. Notably, DOG box dosage was only loosely correlated with expression level of individual effector variants. Comparison of the *G. pallida* genome with those of two other cyst nematodes underlined the fundamental differences in evolutionary history between effector families. Re-sequencing of PCN populations with deviant virulence characteristics will allow for the linking of these characteristics with the composition of the effector repertoire as well as for the mapping of PCN diversification patterns resulting from extreme anthropogenic range expansion.

## 1. INTRODUCTION

Worldwide, affordable food and feed production depends on large-scale monocropping. For practical and economic reasons crop homogeneity in terms of yield quality and quantity is essential. At the same time, such systems are intrinsically vulnerable to damage by pests and pathogens. The highest susceptibility for biotic stressors is found in genetically homogeneous crops. Potato is the third most important staple food (Birch et al., 2012), and in most production systems clonally-propagated seed potatoes are used as starting material. Such production systems cannot do without rigorous disease management. Potato cyst nematodes (PCN), the common name for actually two species, *Globodera pallida*, and *G. rostochiensis*, are among the primary yield-limiting potato pathogens worldwide. PCN co-evolved with potato in the Andes in South America (see e.g. (Plantard et al., 2008)), and proliferation of potato as a main crop outside of its native range was unintentionally paralleled by an enormous range expansion of PCN. For decades, PCN control has mainly been dependent on the application of nematicides. Due to the non-specific nature of these nematicides they have a highly negative impact on the environment, and their use is therefore either banned or severely restricted in most parts of the world. Currently, crop rotation and the use of resistant potato varieties are the main handles in PCN control. For economic reasons, the use of plant resistances is preferred over crop rotation. However, potato resistance genes such as *H1* (Toxopeus & Huijsman, 1952), *Gro1-4* (Paal et al., 2004), *Gpa2* (Bakker et al., 2003), and *H2* (Strachan et al., 2019) are only effective against specific pathotypes of one of these PCN species. Nevertheless, there is no robust (molecular) pathotyping scheme that would allow for the matching of the genetic constitution of field populations with effective host plant resistance genes.

Effectors are proteins secreted by plant-pathogens that facilitate the manipulation of the physiology of the host plant and interfere with the host’s innate immune response in favour of the invading organism (e.g. (Stergiopoulos & De Wit, 2009). Potato cyst nematode effectors have some peculiar characteristics. With at least one known exception, HYPs (Eves-van den Akker, Lilley, Jones, & Urwin, 2014), most effectors are produced in large single-celled glands referred to as the subventral and dorsal esophageal glands. These glands empty into the pharynx lumen, and the lumen is connected to a hollow protrusible stylet with which nematodes pierce plant cell walls. Via the orifice of the stylet, effector proteins are transferred to the apoplast or the cytoplasm of infected host plant cells. It is noted that subventral gland effectors are functional during plant penetration. Subsequently, dorsal gland secretions are responsible for feeding site induction and the suppression of the host’s innate immune system (Smant, Helder, & Goverse, 2018). It has been hypothesised that common transcription factors and/or common promoter motifs might facilitate coordinated expression of effectors during the infection process. Such mechanisms have been identified to regulate effector expression in plant pathogenic fungi and oomycetes (Jones et al., 2019; Roy, Kagda, & Judelson, 2013). Also, among plant-parasitic nematodes, promotor motifs have been identified upstream of effectors that could contribute to the orchestration of the infection process. In the case of the potato cyst nematode *G. rostochiensis*, a DOrsal Gland motif (‘DOG box’) was identified by (Eves-van den Akker et al., 2016). For the pinewood nematode *Bursaphelenchus xylophilus*, a regulatory promotor motif referred to as STATAWAARS was demonstrated to affect effector expression (Espada et al., 2018). Expression of several effectors of Clade I tropical root-knot nematodes (Tandingan De Ley et al., 2002) was suggested to be steered by a putative cis-regulatory motif ‘Mel-DOG’ (*Meloidogyne* DOrsal Gland, (Da Rocha et al., 2021)).

Most likely as a reflection of the co-evolution between nematodes and their host(s), effectors are typically encoded by multigene families showing family-specific levels of diversification (Masonbrink et al., 2019; Van Steenbrugge et al., 2021). Cyst nematodes harbour numerous effector families (see for instance (Pogorelko, Wang, Juvale, Mitchum, & Baum, 2020), and genome (re-)sequencing is a rigorous approach to generate comprehensive overview of PCN effector family compositions. The first genomes of *G. pallida* and *G. rostochiensis* were published by (Cotton et al., 2014) and (Eves-van den Akker et al., 2016). Although this constituted a major step forward, both genomes are very fragmented, hampering effector family inventories. Recently, long-read technology allowed for the generation of a less fragmented and more complete reference genome for *G. rostochiensis* with - among other things - a 24 folds reduction of the number of scaffolds as compared to the initial reference genome (Van Steenbrugge et al., 2021). Here, we present a novel reference genome for the other potato cyst nematode, *G. pallida*, characterised by a 42-fold reduction of the number of scaffolds, together with a reference genome of the beet cyst nematode *Heterodera schachtii*. The *H. schachtii* genome was used to establish the polarity of effectorome contrasts between the two potato cyst nematode species. Detailed knowledge about the nematode’s effector repertoire, a complete overview of variants within effector families, and insights in the evolutionary history of individual effector families are an essential ingredients for a molecular pathotyping scheme. Next to comparing effector diversification patterns, we investigated DOG box distribution and DOG box dosage (up to 16 DOG boxes were observed per putative promoter region) both within and among effectors families. Scrutinising putative effector promoter regions in three reference genomes allowed us to pinpoint the distribution of this putative regulatory motif among cyst nematode species, as well as among and within effector families. Subsequently, the impact of these new, long read technology-based reference genomes on ecological PCN diversification in general and on the development of effectorome-based pathotyping system for potato cyst nematodes in particular is discussed.

## 2. MATERIALS AND METHODS

### 2.1 DNA isolation and sequencing

Cysts from *G. pallida* line D383 were used as starting material for the collection of pre-parasitic second-stage juveniles (J2). J2’s were concentrated, and sucrose centrifugation was used to purify the nematode suspension (Jenkins, 1964). After multiple rounds of washing the purified nematode suspension in 0.1 M NaCl, nematodes were resuspended in sterilised MQ water. Juveniles were lysed in a standard nematode lysis buffer with proteinase K and beta-mercaptoethanol at 60°C for 1 h as described in (Holterman et al., 2006). The lysate was mixed with an equal volume of phenol: chloroform: isoamyl alcohol (25:24:1) (pH 8.0) following a standard DNA purification procedure, and finally, DNA was precipitated with isopropanol. After washing the DNA pellet with 70% ethanol for several times, it was resuspended in 10mM Tris-HCL (pH 8.0). *G. pallida* D383 DNA (10 -20 μg) was sequenced using Pacific Biosciences SMRT sequencing technology at Bioscience (Wageningen Research, Wageningen, The Netherlands). DNA (30 μg) from *H. schachtii* (IRS population) was isolated with a procedure similar to the one used for *G. pallida*, however DNA was precipitated using an ice-cold ethanol precipitation step (Jain et al., 2018). DNA fragments below 10 kb were depleted using a short read eliminator kit (Circulomics SS-100-121-01) and *H. schachtii* DNA (15 μg) was sequenced using Oxford Nanopore technology at NexOmics (Wuhan, China).

### 2.2 Genome assemblies and synteny

For *G. pallida* D383, raw PacBio reads, and for *H. schachtii* IRS, Oxford Nanopore reads were corrected to, in essence, merge haplotypes using the correction mode in Canu (Koren et al., 2017), by reducing the error rate to a maximum of 15% and the corrected coverage to a minimum of 200. Using wtdgb2 v2.3 (Ruan and Li, 2020), multiple initial genome assemblies were generated based on the corrected Nanopore reads while manually refining the parameters minimal read length, k-mer size, and minimal read depth. These parameters were optimised to generate an assembly close to the expected genome size of *G. pallida* and *H. schachtii*. After optimisation, for *G. pallida*, a minimum read length cut-off of 5,000, minimal read depth of 6, and a k-mer size of 18 was used. To generate the assembly of *H. schachtii*, a minimum read length cut-off of 6,000, minimal read depth of 8, and a k-mer size of 21 were used. The remaining haplotigs were pruned from the assemblies using Purge Haplotigs v1.0.4 (Roach *et al*., 2018). The contigs from the assemblies were then improved using FinisherSC v2.1 (Lam *et al*., 2015) at default settings and scaffolded using SSPACE-Longread v1.1 (Boetzer and Pirovano, 2014). Gaps in the assemblies were then filled using GapFiller v1.0 (van Steenbrugge, 2021). For *G. pallida*, the resulting assembly was polished with Pacbio reads by three iterations of Arrow v2.3.3 (https://github.com/PacificBiosciences/GenomicConsensus), followed by five iterations of polishing with Pilon v1.23 (Walker *et al*., 2014) using Pacbio and Illumina NovaSeq reads. Finally, the assembly of *H. schachtii* was polished with Medaka v1.4.1 model r941_prom_hac_g3210 using Nanopore sequencing reads. Repeat regions were softmasked using RepeatModeler v1.0.11 (https://github.com/Dfam-consortium/RepeatModeler) and RepeatMasker v4.0.9 (Tarailo-Graovac and Chen, 2009). Using Braker v2.1.2 (Brůna *et al*., 2021), gene annotations were predicted for both assemblies at default settings outputting gff3 annotations and aided by RNAseq data of different life stages (*G. pallida*: NCBI Bioproject PRJEB2896, *H. schachtii*: PRJNA767548). Full details on the generation of the genome assemblies and prediction of genes are available on https://github.com/Jorisvansteenbrugge/Gros_Gpal_Hsch. For *Globodera rostochiensis*, the Gr-Line19 genome assembly described in (Van Steenbrugge *et al*., 2021) was used (NCBI GenBank assembly accession: GCA_018350325.1).

The synteny between the *G. rostochiensis, G. pallida*, and *H. schachtii* genomes was assessed by a progressive genome alignment using Mauve v2.4.0 (Darling *et al*., 2004). The resulting alignments of regions larger than 1 kb and larger than 3kb were visualised in Circos v0.69-9 (Krzywinski *et al*., 2009).

### 2.3 Identification of Effector Homologs

Effector gene families were identified in the genomes based on the predicted genes by BRAKER2. Homologs for Gr-1106/Hg-GLAND4 were identified using HMMer v3 with a custom HMM profile based on GenBank entries JQ912480 to JQ912513. SPRY homologs were identified with HMMer using a pre-calculated profile HMM in the PFAM database (PF00622). Homologs to CLE-like proteins were identified with a custom profile HMM-based on UniProt sequences (D1FNJ7, D1FNK5, D1FNJ9, D1FNK2, D1FNK8, D1FNK3, D1FNK0, D1FNK4). Homologs to Venom Allergen-like proteins (VAP) were identified with a custom profile HMM-based on Uniprot sequences (Q8MQ79, A0A0K3AST9, P90958, Q19348, A0A0K3AWG2, Q967G4, Q9BID5, A0A3Q8UEU8, Q963I7, B8LF85, A7X975, A0A7G7LJV8). Hg-GLAND5 and Hg-GLAND13 were identified with BLASTP searches with GenBank sequences KJ825716 and KJ825724, respectively, maintaining thresholds at 35% identity, 50% query coverage, and an E-value of 0.0001. Each effector homolog was tested for the presence of a signal peptide for secretion by Phobius v1.01 (Käll, Krogh, & Sonnhammer, 2007) and the presence of one or multiple DOG-box motifs in the promoter region using a custom script (available on GitHub https://github.com/Jorisvansteenbrugge/Gros_Gpal_Hsch).

For *G. rostochiensis*, effector annotations were used as described in (J. J. M. van Steenbrugge et al., 2021), except for CLE-like proteins. CLE variant Gros19_g16105.t1 was excluded since the gene model likely contains errors, and the exact location of this variant in the phylogenetic tree is therefore uncertain. Furthermore, the HMM scoring cut-off was lowered to 300 to include two more potential CLE variants.

### 2.4 Phylogeny

A multiple sequence alignment was generated for each effector gene family using Muscle v3.8.1551 based on gene coding sequences to infer the phylogeny of effector gene families between species. Next, phylogenetic trees were produced with RaxML v8.2.12 (Stamatakis, 2014) using the GTRGAMMA model with 100 bootstrap replicates. The GTRGAMMA model was selected based on recommendations by ModelTest-NG v0.2.0 (Darriba et al., 2020). Finally, using Figtree V1.4.4, the resulting trees were visualised and annotated.

### 2.5 DOGbox identification and Gene expression Analysis

RNAseq reads (add repository) were mapped to the *G. pallida* D383 genome, Reads xxx were mapped to the *G. rostochiensis* Gr-Line19 genome, and Reads xxx were mapped to the *H. schachtii* IRS genome using Hisat2 v2.1.0, generating alignments tailored for transcript assemblers (--dta option). Based on the alignments, abundances were estimated and normalized to transcripts per million (TPM) for the transcripts predicted by Braker2 using StringTie v2.1.7b (Kovaka et al., 2019). Expression values of SPRYSEC genes in all three cyst nematode species were extracted, and plotted against the number of DOG-box motifs in each gene. Spearman’s rank-order correlation was used to determine relationship between TPM and the number of DOG-box motifs.

## 3. RESULTS

### 3.1. The use of long read sequence technologies to generate novel reference genomes

The mapping of diversification patterns of effector families requires a high-quality reference genome with preferably a low number of scaffolds and a minimal gap length. (Cotton et al., 2014) were the first to present a reference genome of the potato cyst nematode species *G. pallida*. For our specific purpose, *i*.*e*., the generation of complete inventories of effector families, this reference genome was too fragmented, and the gap length was too large. PacBio long-read technology allowed us to generate a new reference genome from the *G. pallida* population D383 with a 42-fold reduction of scaffolds and a 21-fold reduction of the total gap length. As one of the results, the number of predicted genes increased from 16,403 to 18,813.

In addition, we assembled the genome sequence of the IRS population of beet cyst nematode *Heterodera schachtii*. This allowed us to compare effector family diversification among the two potato cyst nematode species, *G. pallida* and *G. rostochiensis*, and establish the polarity of these contrasts by using *H. schachtii* as an outgroup (both *Globodera* and *Heterodera* belong to the family Heteroderidae). The current genome size, 190 Mb, is slightly above the genome size estimated by flow cytometry, 160 - 170 Mb (Eves van den Akker, personal communication). It is noted that both the predicted number of genes and transcripts were about 50% higher in *H. schachtii* than in the two *Globodera species*.

Two synteny plots were generated based on the alignment of regions >1 kb and >3kb to compare the genomic organisation of the three cyst nematode species. Not unexpectedly, the two potato cyst nematode species share numerous >1 kb regions (Fig. 1A). In the *H. schachtii* genome, several homologous >1kb regions cluster together in genomic segments that span over 2 Mb (Fig. 1A, segments 1-8). It is noted that the homologous >1kb regions in these segments have equivalents in both *G. pallida* and *G. rostochiensis*. Alignment of >3kb fragments severely reduced the number of homologous regions among the three cyst nematode species (Fig. 1B). Nevertheless, the number of shared >3kb regions between *G. pallida* and *G. rostochiensis* (N: 76) is considerably higher than the number of shared regions between *H. schachtii* and *G. rostochiensis* (N: 23) (Fig. 1B).

**Fig. 1.**
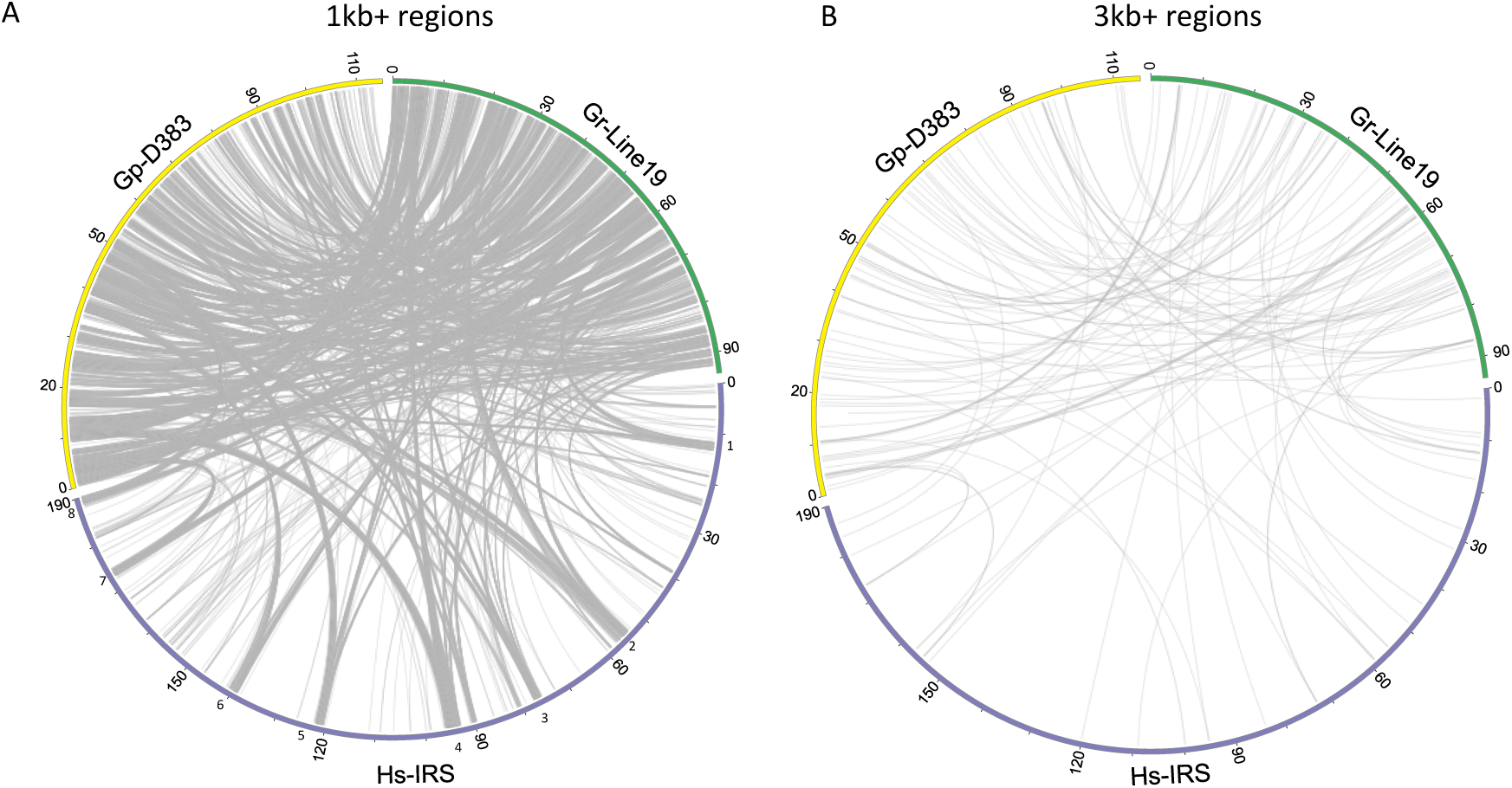
Synteny between *G. pallida* (population D383), *G. rostochiensis* (Gr-Line19) and *H. schachtii* (population IRS) based on a progressive genome alignment in Mauve. In Fig. 1 A syntenic regions larger than 1 kb are shown, in panel B syntenic regions larger than 3 kb are shown. In panel A, *H. schachtii* genome regions are indicated where multiple syntenic regions cluster together into segments spanning over 2 Mb (Fig. 1A, segments 1-8). It is noted that these segments have equivalents in both *G. pallida* and *G. rostochiensis*.

### 3.2 Effector family selection

In our comparison between the three cyst nematode species, we concentrated on effectors. Cyst nematodes were shown to harbour numerous effectors families (Pogorelko et al., 2020). Here we concentrated on six effector families. For four of these families, one or more representatives are known to be involved in the suppression of plant innate immune system (SPRYSEC (Diaz-Granados, Petrescu, Goverse, & Smant, 2016; Mei et al., 2018), GLAND4 (also referred to as Gr-1106) (Barnes, Wram, Mitchum, & Baum, 2018), GLAND5 (also referred to as G11A06) (Yang et al., 2019), and VAP (Wilbers et al., 2018)). CLE (Wang et al., 2021) is an intriguing effector family involved in feeding site induction, and the GLAND13 (Danchin, Guzeeva, Mantelin, Berepiki, & Jones, 2016) family is essential in the hydrolysis of plant sugars once they are taken up by the nematode.

### 3.3. SPRYSECs

SPRYSEC is an acronym for a family of secreted effectors with an SP1a/RYanodine receptor domain. This family was recently shown to be highly diverged in the potato cyst nematode *G. rostochiensis* (J. J. M. van Steenbrugge et al., 2021). Fig. 2 shows a phylogenetic tree with (supposedly) all SPRYSECs present in the three cyst nematode species under investigation. The number of paralogs in *G. pallida, G. rostochiensis*, and *H. schachtii* is respectively 24, 60 and 13. Despite the poor backbone resolution of the SPRYSEC tree, two moderately supported SPRYSEC clades (A and B) could be distinguished. Clade A comprises SPRYSEC variants exclusively from the two potato cyst nematode species, and *G. pallida* SPRYSEC paralogs are interspersed with *G. rostochiensis* SPRYSEC variants. SPRYSEC Clade A is characterised by a dosage of 0 – 6 DOG box elements. Clade B harbours fewer SPRYSEC paralogs than Clade A (27 *versus* 35 in Clade A). Notably, Gr19_g7942 is not preceded by a signal peptide for secretion, whereas three DOG box elements are present in the promoter region directly upstream of this paralog. Clade B is characterised by a mix of SPRYSEC variants solely originating from *G. pallida* and *G. rostochiensis*. Compared to Clade A, Clade B is typified by an overall higher DOG box dosage (on average, 1.7 and 5.2 DOG boxes per paralog). Up to 16 DOG box elements were identified in the promoter regions of paralogs in Clade B.

**Fig. 2.**
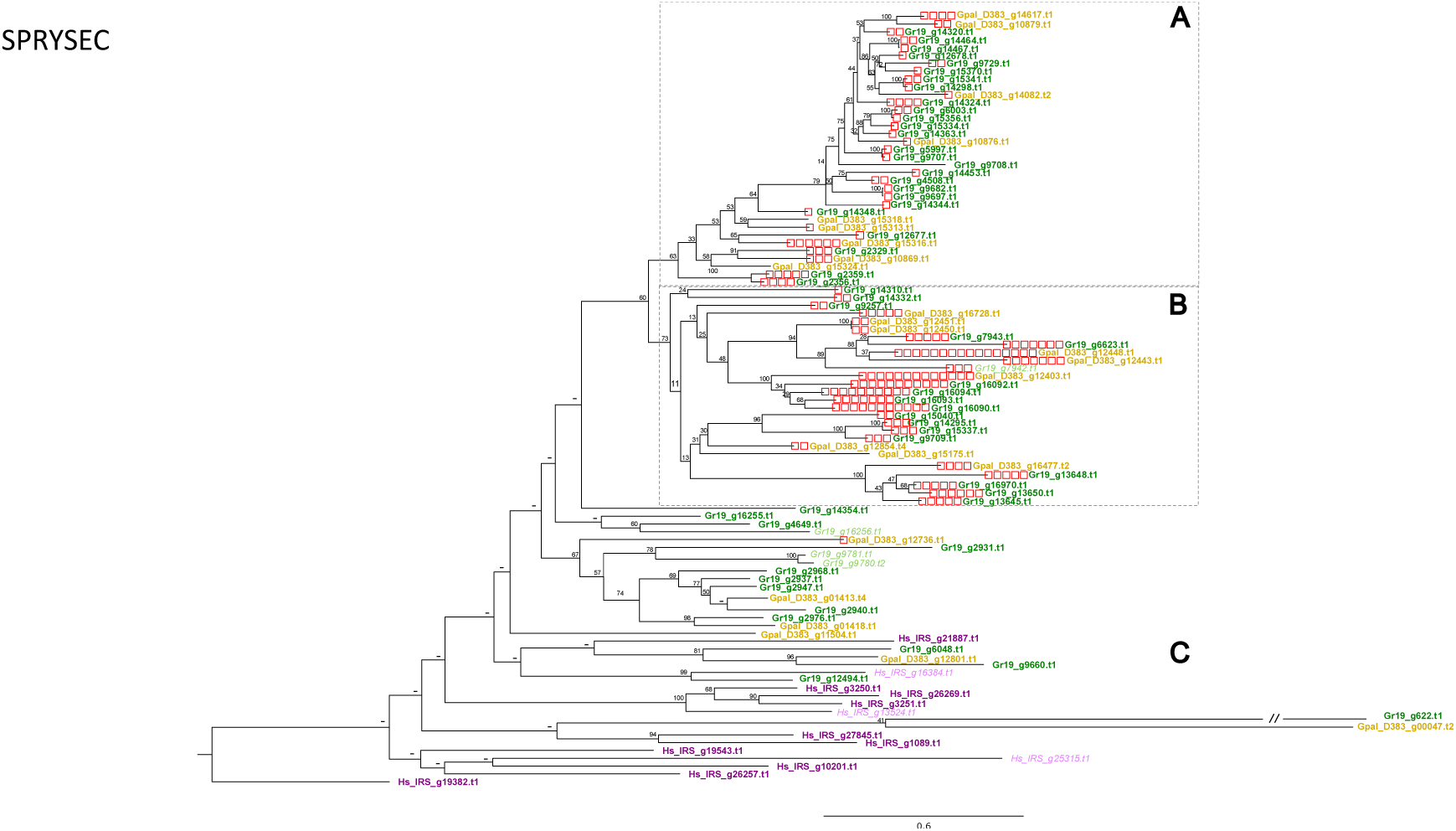
Phylogeny of SPRYSEC effector genes (see e.g. (Diaz-Granados et al., 2016)) of *G. pallida* (population D383) (ochre), *G. rostochiensis* (Gr-Line19) (green) and *H. schachtii* (population IRS) (purple). A multiple sequence alignment was made using MUSCLE on the coding sequence. A phylogenetic tree was made using RAxML using a GTRGAMMA model, validated by 100 bootstrap replicates. Bootstrap values < 50 % are indicated by a “-”. Gen IDs in italics in lighter shades of ochre, green or purple are used to indicate effector variants with at least one predicted transmembrane domain. Boxed clusters (A and B) highlight two moderated supported subclades with on average moderate (A) and high DOG box dosages. C refers to the basal part of the SPRYSEC tree.

The more basal part of the SPRYSEC tree (Fig. 2, part C) harbours, next to paralogs from the two potato cyst nematode species, all 13 SPRYSEC variants from *H. schachtii*. None of the promoter regions of the 35 SPRYSEC paralogs in this part of the SPRYSEC tree harbours DOG boxes. Both *H. schachtii* and *G. rostochiensis* harbour SPRYSEC paralogs with at least one transmembrane domain (gene ID in italics with lighter colour).

### 3.4. GLAND4

The number of GLAND4 (also referred to as Gr-1106) paralogs in Gr-Line19, Gp-D383, and Hs-IRS is respectively 10, 9, and 15. The phylogenetic analysis yields a tree with a well-supported backbone (Fig. 3) showing a clear separation between the outgroup *H. schachtii*, and both *Globodera* species. While the GLAND4 paralogs in Gr-Line19 and Gp-D383 show little divergence, they end up in separate species-specific clusters. On the other hand, Hs-IRS paralogs show more intraspecific diversification. All but two *G. pallida* paralogs (Gpal_D383_g17346.t1 and Gpal_D383_g13669) contain a signal peptide for secretion. For all but one of the GLAND4 genes in Gr-Line19, the promoter region included a DOG-box motif, while promoter regions of only four GLAND4 genes in Gp-D383 and one in Hs-IRS contained such a motif.

**Fig. 3.**
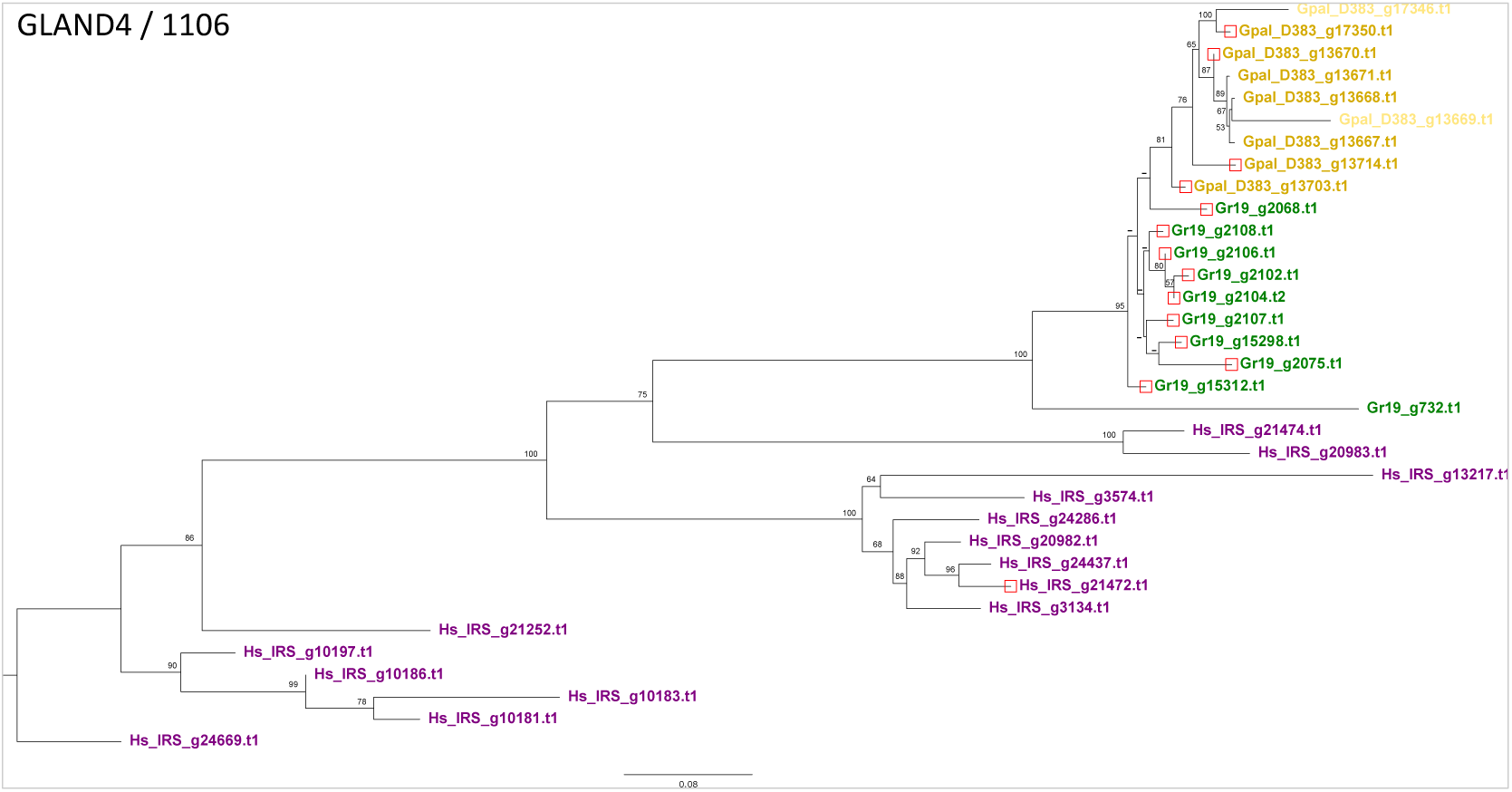
Phylogeny of GLAND 4 (equivalent to 1106, see, e.g., (Noon et al., 2015)) effector genes of *G. pallida* (population D383) (ochre), *G. rostochiensis* (Gr-Line19) (green) and *H. schachtii* (population IRS) (purple). A multiple sequence alignment was made using MUSCLE on the coding sequence. A phylogenetic tree was made using RAxML using a GTRGAMMA model, validated by 100 bootstrap replicates. Bootstrap values < 50 % are indicated by a “-”. Bootstrap values < 50 % are indicated by a “-”. Gen IDs in lighter shades of ochre, green or purple are used to indicate effector variants lacking a signal peptide for secretion.

### 3.5. GLAND5

With 13 homologs, the GLAND5 effector family (also referred to as G11A06) was significantly less diversified in Gr-Line19 than in Gp-D383 and Hs-IRS with respectively 25 and 27 paralogs. In all three species, the majority of the GLAND5 paralogs harbour a signal peptide for secretion. It is worth noting that a relatively high percentage of the GLAND5 paralogs in Gr-Line19 was not preceded by a signal peptide for secretion (23%). In contrast, in *H. schachtii* and *G. pallida*, respectively, 88.9% and 92% of the paralogs comprised a signal peptide. The phylogenetic analysis (Fig. 4) shows that GLAND5 is a diversified gene family. Several well-supported branching events define a set of subclades that either exclusively comprises *H. schachtii* or contain GLAND5 variants from both potato cyst nematode species in the more distal branches. Even though the GLAND5 paralogs Gr-Line19 and Gp-D383 occur together in individual subclades, no obvious sets of potential orthologs between the two species could be identified. In *H. schachtii*, 82% of the paralogs contain at least one DOG-box motif in the promoter region. Out of the three GLAND5 paralogs without a signal peptide for secretion, two (Hs-IRS_g6495.t1 and Hs-IRS_g22438.t1) had at least one DOG box motif in their promoter region. DOG boxes were less prominently present among the *G. rostochiensis* and *G. pallida* GLAND5 variants (39% and 20% of the paralogs).

**Fig. 4.**
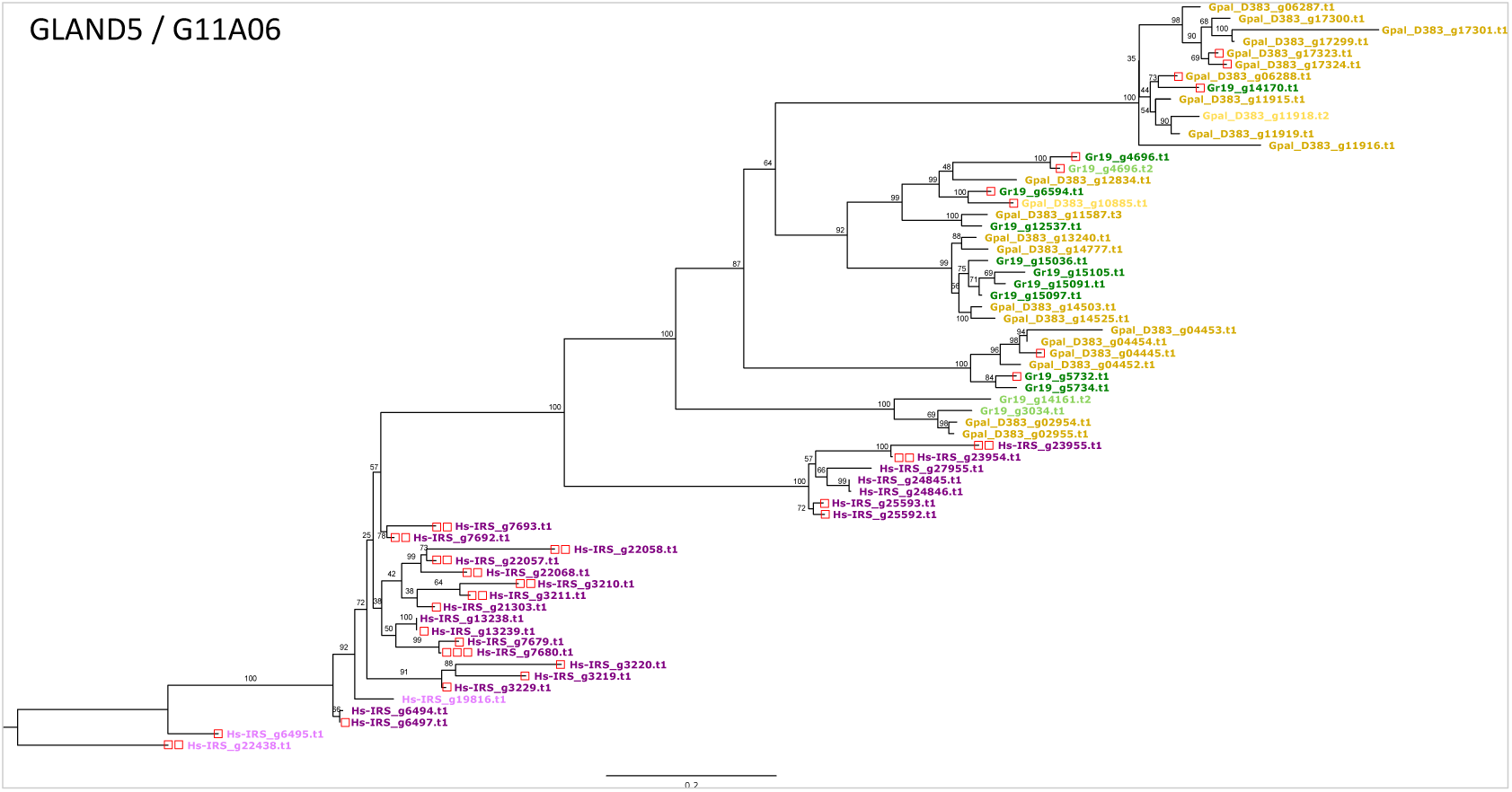
Phylogeny of GLAND 5 (equivalent to G11A06, see, e.g., (Noon et al., 2015)) effector genes of *G. pallida* (population D383) (ochre), *G. rostochiensis* (Gr-Line19) (green) and *H. schachtii* (population IRS) (purple). A multiple sequence alignment was made using MUSCLE on the coding sequence. A phylogenetic tree was made using RAxML using a GTRGAMMA model, validated by 100 bootstrap replicates. Bootstrap values < 50 % are indicated by a “-”. Bootstrap values < 50 % are indicated by a “-”. Gen IDs in lighter shades of ochre, green or purple are used to indicate effector variants lacking a signal peptide for secretion.

### 3.6. VAP (Venom Allergen-like Proteins)

The levels of diversification in the VAP effector family were highly comparable between the three cyst nematode species. In Gp-D383, Hs-IRS, and Gr-Line19, respectively, 8, 8, and 9 VAP paralogs were identified. The phylogenetic analysis resulted in a tree with a well-supported backbone (Fig. 5). The tree contains three clusters (Fig. 5, boxes A, B, C) with a high level of diversification between the clusters. At the base of the tree, a small cluster of 4 *H. schachtii* paralogs (Fig. 6, box C) is present that all lack a signal peptide for secretion. Box B harbours Gp-D383 and Gr-Line19 VAP paralogs, of which all but two lack a signal peptide for secretion. Three sub-clusters are present in Box A: one with exclusively Gr-Line19 variants, a second one with just Gp-D383 variants, and the third with an orthologous pair between Gr-Line19 and Gp-D383. In the largest cluster at the top of the tree (Fig. 5, box A), VAP paralogs of all three species are present, including the only secreted VAP variant in Hs-IRS with a DOG-box motif in the promoter region.

**Fig. 5.**
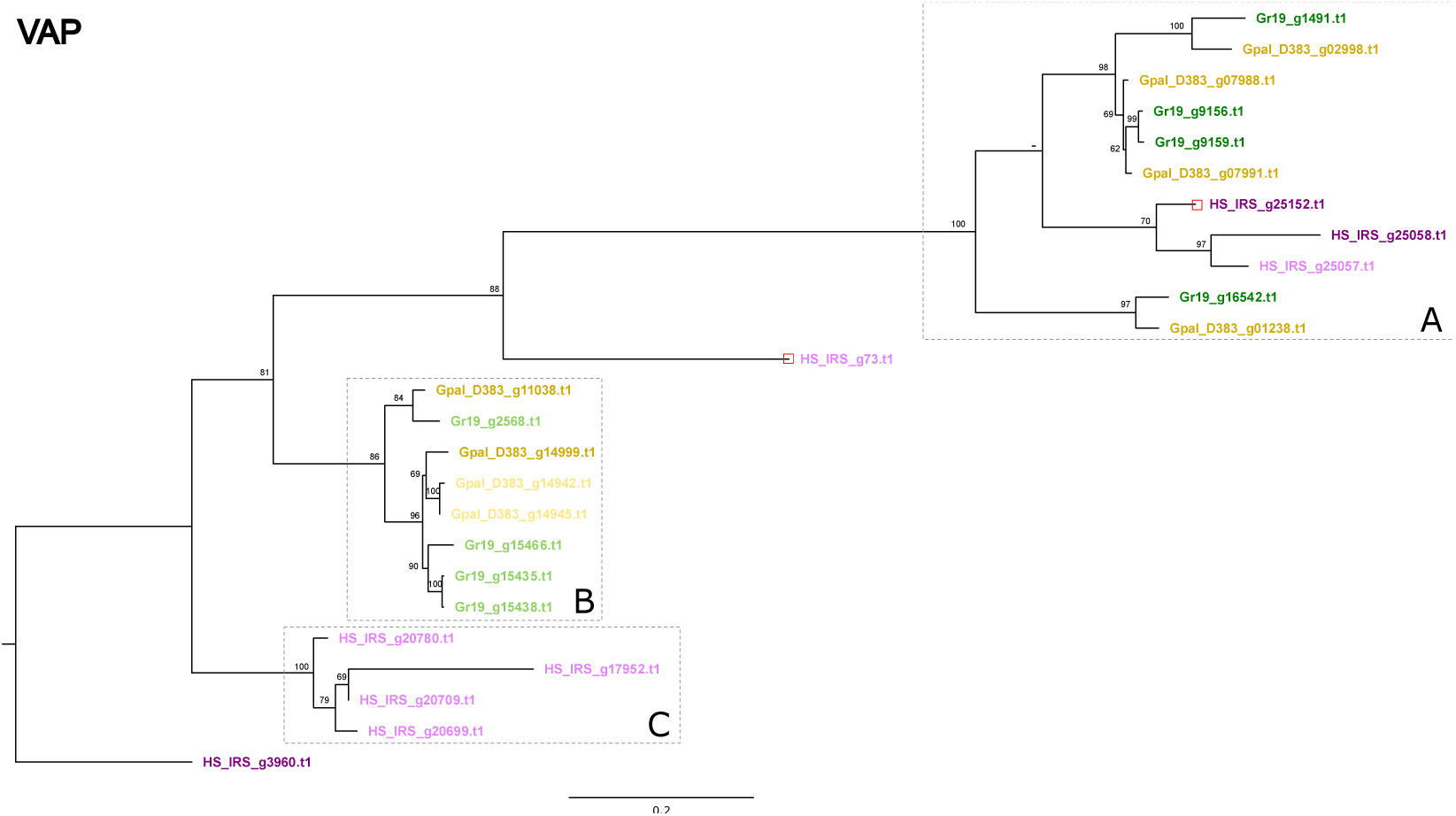
Phylogeny of VAP (Venom Allergen-like Protein, see, e.g., (Wilbers et al., 2018)) effector genes of *G. pallida* (population D383) (ochre), *G. rostochiensis* (Gr-Line19) (green) and *H. schachtii* (population IRS) (purple). A multiple sequence alignment was made using MUSCLE on the coding sequence. A phylogenetic tree was made using RAxML using a GTRGAMMA model, validated by 100 bootstrap replicates. Bootstrap values < 50 % are indicated by a “-”. Bootstrap values < 50 % are indicated by a “-”. Gen IDs in lighter shades of ochre, green or purple are used to indicate effector variants lacking a signal peptide for secretion.

**Fig. 6.**
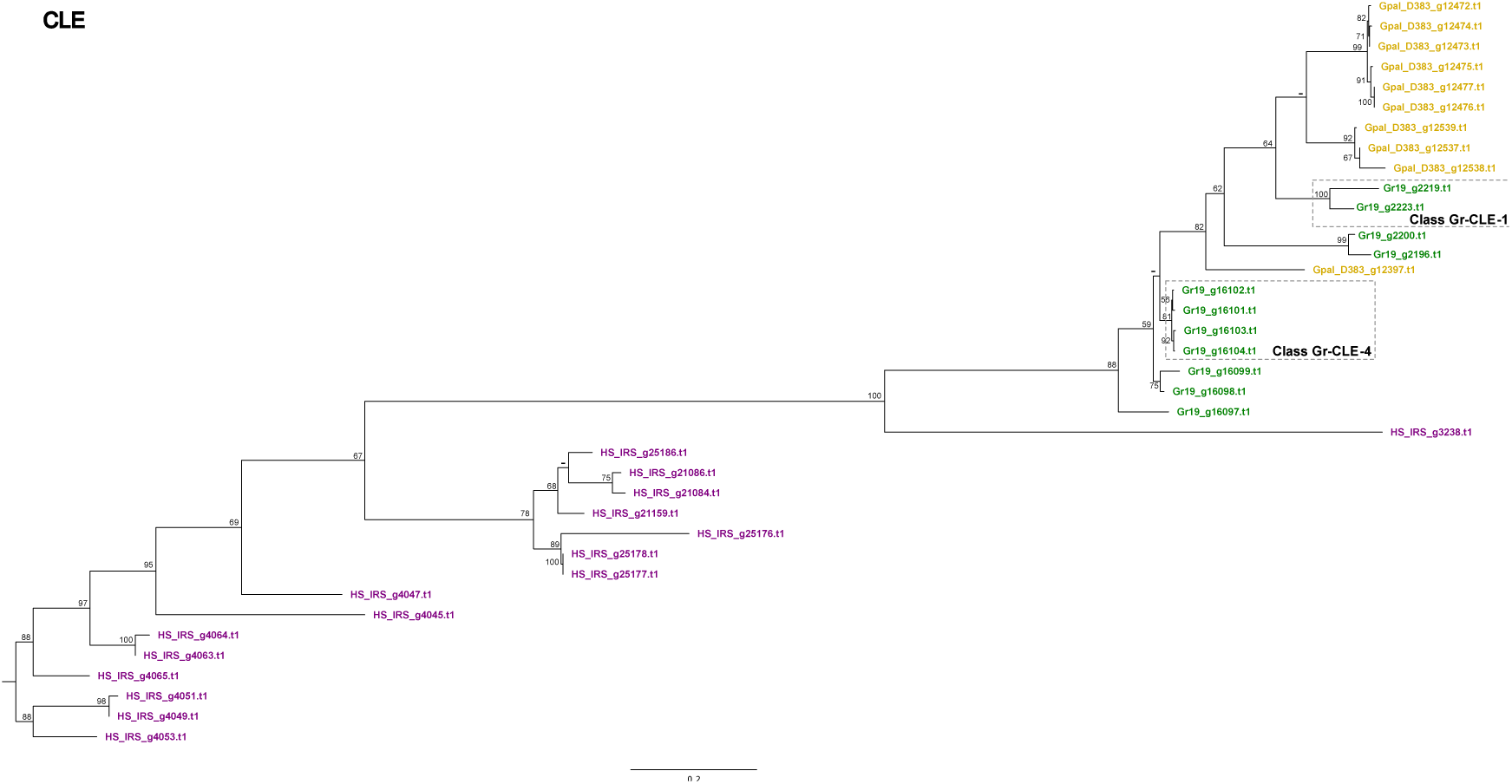
Phylogeny of CLE (CLAVATA3/ESR-related peptides, see, e.g., (Lu et al., 2009)) effector genes of *G. pallida* (population D383) (ochre), *G. rostochiensis* (Gr-Line19) (green) and *H. schachtii* (population IRS) (purple). A multiple sequence alignment was made using MUSCLE on the coding sequence. A phylogenetic tree was made using RAxML using a GTRGAMMA model, validated by 100 bootstrap replicates. Bootstrap values < 50 % are indicated by a “-”. Bootstrap values < 50 % are indicated by a “-”. Gen IDs in lighter shades of ochre, green or purple are used to indicate effector variants lacking a signal peptide for secretion.

### 3.7. CLE (<u>CL</u>AVATA3/<u>E</u>SR-related peptides)

With 16 variants, the CLE-like effector family is considerably more diversified in *H. schachtii* than in *G. pallida* and *G. rostochiensis* (respectively, 10 and 11 paralogs). Analysis of the CLE families on the three cyst nematode species resulted in a phylogenetic tree with a reasonably well-resolved backbone (Fig. 6). It is worth noting that nearly all variants are united in species-specific clusters, and in this sense, the CLE diversification patterns resemble the patterns observed for the GLAND4 (Gr-1106) family (Fig. 3). Whereas Gp-D383 and Gr-Line19 are characterised by similar-sized, moderately diverged clusters of CLE paralogs, the CLE family has much more diverged in *H. schachtii*.

In *G. rostochiensis*, two functional classes of CLE peptides have been described, CLE-1 and CLE-4 (Lu et al. 2009). The CLE-1 class (Fig. 6) comprises two Gr-Line19 paralogs that show only distant homology to *G. pallida* CLEs. Similarly, four *G. rostochiensis* CLE’s belonging to functional CLE class 4 (Fig. 6) do not have clear equivalents in *G. pallida* and *H. schachtii*. Unlike all other effector families investigated so far, all CLE variants from the three cyst nematode species are preceded by a signal peptide for secretion. At the same time, none of them has a DOG-box motif in the promoter region.

### 3.8. GLAND13 (Glycosyl Hydrolase Family 32 (GH32) members)

GLAND13 effectors investigated so far were identified in *G. pallida* and coded for invertases belonging to glycosyl hydrolase family 32 (GH32). While these enzymes are not secreted into the plant, they are essential as they catalyse the hydrolysis of the primary type of sugar the nematode takes up from its host, sucrose (Danchin et al., 2016). This gene family shows a large difference in the number of paralogs present in the three species; while Gr-Line19 and Gp-D383 harbour 10 and 7 paralogs, Hs-IRS holds only 3 copies. In the phylogenetic tree (Fig. 7), the paralogs in *H. schachtii* are positioned at the tree’s base. In Box A, paralogs of Gr-Line19 and Gp-D383 are interspersed, while in Box B, all but one paralogs are from Gr-Line19. In Gr-Line19, 70% of the GLAND13 paralogs comprise a signal peptide for secretion, slightly lower percentages (67% and 57%) were observed in Hs-IRS and Gp-D383. Variants showing high similarity to each of the five GLAND13 paralogs from *G. pallida* (population Lindley; indicated as GPLIN_*number*) are indicated in Fig. 7.

**Fig. 7.**
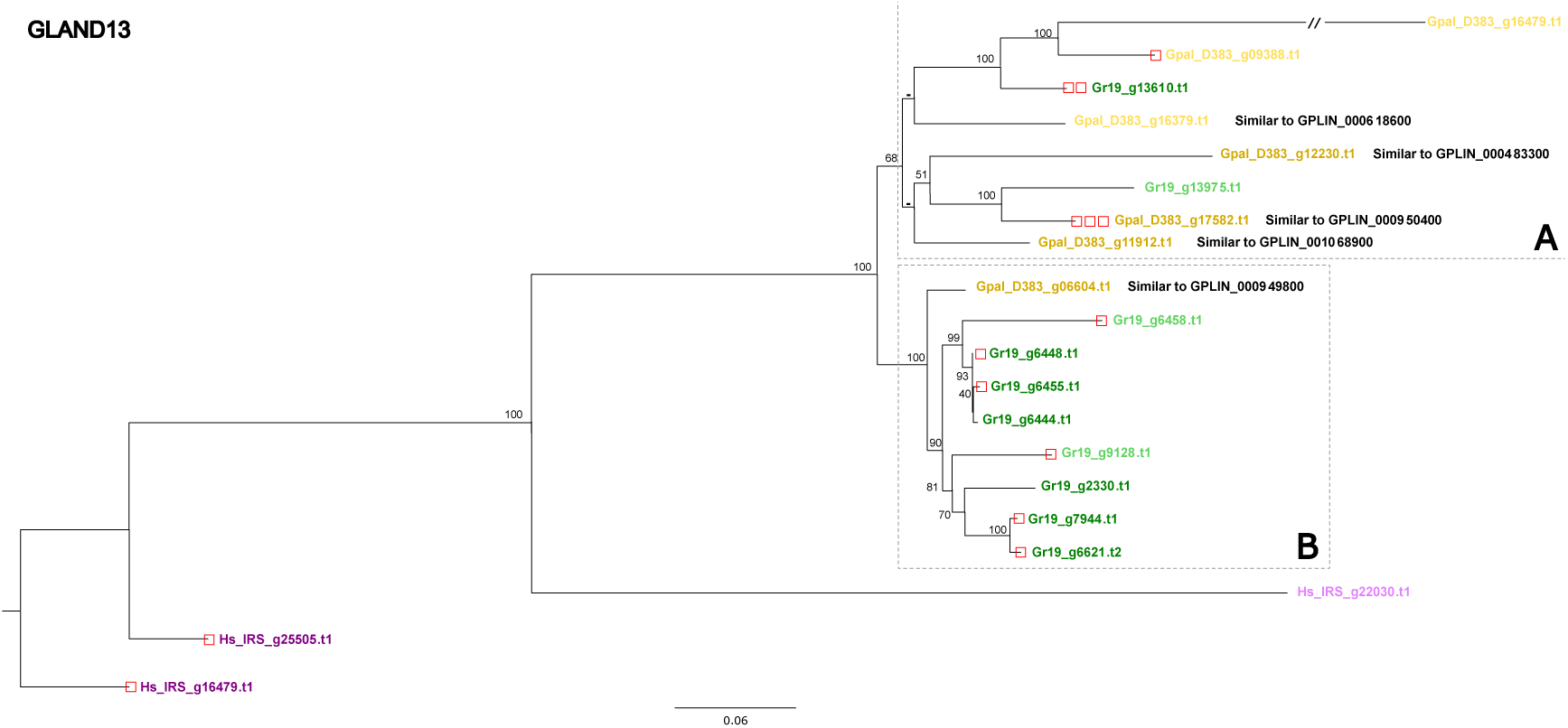
Phylogeny of GLAND13 (invertases, see, e.g., (Danchin et al., 2016)) effector genes of *G. pallida* (population D383) (ochre), *G. rostochiensis* (Gr-Line19) (green) and *H. schachtii* (population IRS) (purple). A multiple sequence alignment was made using MUSCLE on the coding sequence. A phylogenetic tree was made using RAxML using a GTRGAMMA model, validated by 100 bootstrap replicates. Bootstrap values < 50 % are indicated by a “-”. Bootstrap values < 50 % are indicated by a “-”. Gen IDs in lighter shades of ochre, green or purple are used to indicate effector variants lacking a signal peptide for secretion.

For half of the GLAND13 effector variants in Gr-Line19, a DOG-box motif in the promoter region was shown. One gene, Gr19_g13610, contained the motif two times. In *H. schachtii*, this motif was present in 2 out of 3 genes, while in *G. pallida*, DOG boxes were found in one variant with a signal peptide for secretion (Gpal_D383_g17582), and in one paralog without such a signal (Gpal_D383_g09388). It is noted that these DOG box motifs were found in promoter regions of *G. pallida* effectors that are not expressed in the dorsal gland (Danchin et al., 2016).

### 3.9 Effects of DOG-box dosage on SPRYSEC Expression

Although DOG-box motifs in the promoter regions of effector variants are present in many effector families, their presence is not a necessity for the functioning of an effector family. For example, none of the variants of the CLE family contained DOG-box motifs, regardless of the species (Fig. 6). Dorsal gland-expressed effectors can therefore be expressed and secreted without the presence of DOG-box motifs. This is further illustrated in Fig. 8A, this bar chart shows DOG box distribution of over the six effector families. For the three cyst nematode species under investigation it demonstrates that DOG boxes can be entirely absent (CLE), present in some species only (VAP, SPRYSEC), or present in all species (GLAND4, GLAND5, GLAND13). The ample presence of DOG boxes in the diversified SPRYSEC family prompted us to investigate whether there is a correlation between the DOG-box dosage and SPRYSEC expression levels. SPRYSEC effectors from all three cyst nematode species were taking into account (Fig. 8B). A modest correlation (R^2^= 0.66) between the DOG-box dosage and the expression levels based on RNA abundance is present for this family. Especially in *G. pallida* high expression levels of SPRYSEC variants can be reached in absence of DOG boxes in its promoter region (Fig. 8B).

**Fig. 8.**
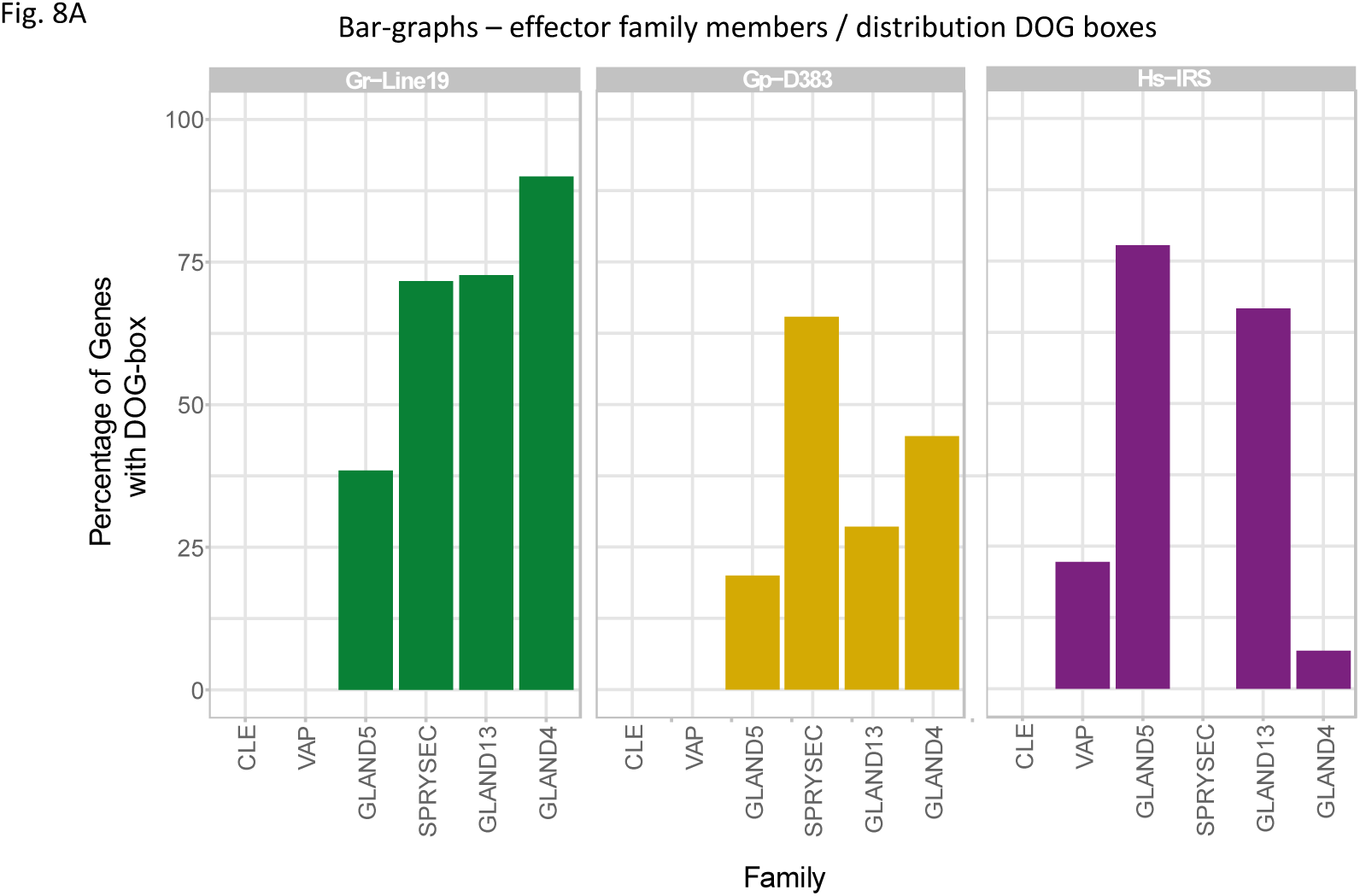
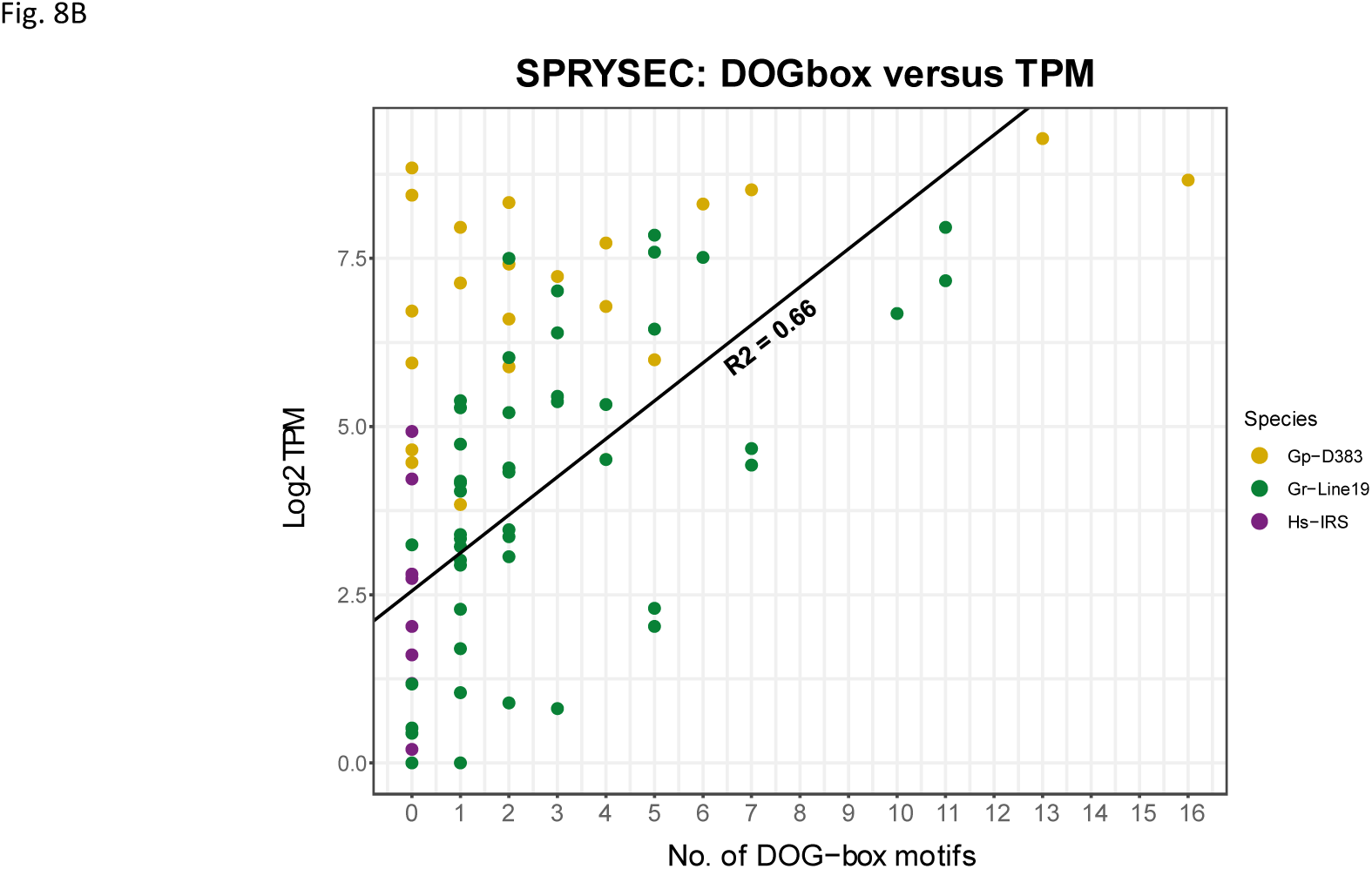
This distribution of a DOrsal Gland promoter element motif (‘DOG box, (Eves-van den Akker et al., 2016)) among a selection of cyst nematode species. Fig. 8A shows the percentages of variants per effector family with one or more a DOG boxes in their promoter region. In Fig. 8B the relationship between DOG box dosage and expression level (expressed as log2 TPM (Transcript Count Per Million) is presented.

## 4. DISCUSSION

In our attempts to fundamentally understand the interaction between plant-parasitic nematodes and their hosts, the usefulness of high-quality reference genomes of these pathogens is beyond discussion. Keeping in mind the enormous impact of PCN in all major potato production regions of the world, it is not surprising that high priority was given to the sequencing of both the *G. pallida* (Cotton et al., 2014) and the *G. rostochiensis* (Eves-van den Akker et al., 2016) genome. This was done before long-read sequencing technologies became available. Although some research questions can very well be addressed with these reference genomes, less fragmented genomes are needed for studying effector diversification. Therefore, a new reference genome was generated from *G. pallida* population D383. As compared to the *G. pallida* Lindley genome assembly, this resulted in a 42-fold reduction in the number of scaffolds and a 24-times increase of the N50. In the comparison of the effectoromes of the two PCN species, we included a newly generated genome of the H. schachtii population IRS as an outgroup. It is noted that reference genomes from these obligatory sexually reproducing pathogens are actually population-based consensus genomes. Long read sequencing technologies require DNA from 10,000s genetically non-identical nematodes. While an individual of these diploid species could theoretically carry a maximum of two haplotypes per locus, a population has the potential to carry many more. It is essential to mine these haplotypes and assemble them into a single haploid assembly to generate a proper reference. This is not a trivial process and requires specialised bioinformatics software (Roach et al., 2018). As the sizes of the current genome assemblies are comparable to the genome sizes assessed by flow cytometry, and as the BUSCO duplication scores are relatively low, we assume that the current genomes assemblies are a reasonable reflection of their actual constitution.

### 4.1 Effector diversification

In our analyses we concentrated on six selected effector families, and this selection included relatively widespread effector families such as the CLE, GLAND13 and the VAP family, as well as families that appear to be cyst nematode lineage specific such as SPRYSEC, GLAND4 and GLAND5. Although the protein architecture is distinct between lineages (see e.g. (Mitchum, Wang, Wang, & Davis, 2012)), the CLE family - a category of effectors involved in feeding site induction - were shown to be present as well in root-knot and reniform nematodes (Rutter et al., 2014; Wubben, Gavilano, Baum, & Davis, 2015). GLAND13 effectors, members of glycosyl hydrolase family 32, were shown to be present in a range of root knot and cyst nematodes species as well as in other plant-parasitic nematodes such as *Nacobbus aberrans* and *Rotylenchus reniformis* (Danchin et al., 2016). The distribution of VAP within the phylum Nematoda is even broader (Wilbers et al., 2018). Venom allergen-like proteins were discovered in the animal parasite *Ancylostoma caninum* (Hawdon, Jones, Hoffman, & Hotez, 1996). Later on it was isolated from the root-knot nematode *Meloidogyne incognita* (Ding, Shields, Allen, & Hussey, 2000), and subsequently in a wide range of obligatory plant parasitic nematode including various cyst nematode species. A number VAP variants were shown to be implicated in the suppression of both PAMP triggered immunity and effector triggered immunity (e.g. (Li et al., 2021) for the burrowing nematode *Radopholus similis*). Our effector family selection also included families that (so far) appear to be specific to the cyst nematode lineages. This includes SPRYSEC (Diaz-Granados et al., 2016), GLAND4 (also referred to as Gr-1106), and GLAND5 (also referred to as G11A06). For all of these effector families it can be said that at least a subset was shown to be involved in repression of the host plant immune system.

While comparing the overall diversification patterns of the six effector families under investigations, striking differences are observed. In case of SPRYSEC, GLAND5 (G11A06), and GLAND13 (GH32 members), virtually all *H. schachtii* paralogs appeared to be phylogenetically isolated from the *G. pallida* and *G. rostochiensis* effector family variants, while representatives from the two potato cyst nematode species were presented in mixed clusters. These results should be taken with some caution as the backbone resolution of these phylogenetic trees ranges from poor (SPRYSEC) to robust (GLAND5, GLAND13). These patterns suggest that SPRYSEC, GLAND5 and GLAND13 effectors started to diversify after the split between *Heterodera* and *Globodera*.

Effector families GLAND4 (Gr-1106) and CLE showed distinct diversification patterns; by far most paralogs are grouped in species-specific clusters. Keeping in mind that both effector families show a reasonable backbone resolution, we hypothesize that these effector families might have diverged after the split between *G. pallida* and *G. rostochiensis*.

Phylogenetic analysis of the VAP effector family in the three cyst nematode species revealed an opposite pattern as an almost complete mixtures of representative paralogs from the individual species was observed. VAPs constitute an exceptionally widespread effector family within the phylum Nematoda (Wilbers et al., 2018), and our results point at diversification of this family before the split between the cyst nematode genera *Globodera* and *Heterodera*.

### 4.2 Regulation of effector gene expression

Various stages of the parasitic life cycle of cyst nematodes such as plant invasion, feeding site induction and feeding site maintenance require the carefully orchestrated expression of distinct blends of effector proteins (Elashry et al., 2020; Thorpe et al., 2014). For some obligatory plant-parasitic nematodes promoter elements have been identified that were suggested to be involved in this orchestration (Da Rocha et al., 2021; Eves-van den Akker et al., 2016). For the three cyst nematode species we showed the presence of a short DNA box motif (‘DOG box’; ATGCCA) in the promoter region of some members of some of the effector families. The absence of DOG boxes in the CLE family, the scattered presence of DOG boxes in the other 5 families and the loose correlation between DOG box dosage and expression level, prompt us to conclude that DOG boxes might contribute to the orchestration of effector expression, but we see little evidence for a central role of this DNA motif in this process. Further investigation is necessary to elucidate the function of DOG boxes in effector regulation.

In case of plant pathogenic fungi, a few transcription factor have been identified that were shown to steer effector expression. SIX Gene Expression 1 (Sge1), a conserved member of Gti1/Pac2 protein family, was instrumental in the regulation of effector repression in a range of fungal pathogens including *Verticillium dahlia* (Santhanam & Thomma, 2013), *Zymoseptoria tritici* (Mirzadi Gohari et al., 2014), and *F. oxysporum f. sp. cubense* (Hou et al., 2018). As another example AbPf2 could be mentioned, a zinc cluster transcription factor from the necrotrophic plant pathogen *Alternaria brassicicola*. Via a loss of function approach, this transcription factor was shown to regulate the expression of eight putative effectors (Cho, Ohm, Grigoriev, & Srivastava, 2013). Evidently, plant pathogenic fungi are only very distantly related to plant parasitic nematodes, and these examples should only be considered as an illustration how effector expression is organized in other pant pathogen systems.

## 5. CONCLUSIONS

The potato cyst nematode *Globodera pallida* and its sibling species *G. rostochiensis* coevolved with potato in the Andes in South America. These pathogens have been introduced unintentionally in all major potato growing regions in the world. Currently, PCN belongs to the most harmful pathogens in potato production systems, and as a result of this extreme anthropogenic range expansion potatoes worldwide can’t be grown without adequate PCN management. For both *G. pallida* and *G. rostochiensis* host plant resistances belong currently to most powerful means to control these soil pathogens. However, their effectiveness depends on the proper matching between the genetic constitution of the PCN field population and the set of host pant resistances present in modern potato varieties. In 2014, (Niere, Krüssel, & Osmers, 2014) reported about *G. pallida* populations that could no longer be controlled by any of the currently used potato cultivars. This, combined with inherent imperfectness of the current *G. pallida* pathotyping system (e.g. (Phillips & Trudgill, 1983)), underlines the need for a new pathotyping system. Such a system will be based on distinctive effector variations present in any given PCN population. The availability of a high quality reference genome is a prerequisite for the development of such a system. We demonstrated that the quality of the *G. pallida* genome presented in this paper allows for the mapping of complete effector families. With this resource, re-sequencing data from pathotypically diverse *G. pallida* populations will provide insight in the ecological diversification of this extreme range expander, and enable the development of a new pathotyping system that will facilitate the targeted and durable use of precious host plant resistances against this notorious plant pathogen.

## ACKNOWLEDGEMENTS

JvS, MH and SvdE were supported by a grant from the Applied and Technical Science domain (TTW) of the Netherlands Organization for Scientific Research (NWO) under grant no. 14708. PT received support from the University of St Andrews Bioinformatics Unit (AMD3BIOINF), funded by Wellcome Trust ISSF award 105621/Z/14/Z. MS benefitted from funding by a VENI grant (17282) from the NWO domain Applied and Engineering Sciences

## AUTHOR CONTRIBUTIONS

SvdE performed the DNA extraction and library preparations for PCN. JL performed the DNA extraction and library preparations for *H. schachtii*. JvS, PT, and MH conceptualised the genome assembly pipeline. JvS and MH generated the genome assemblies. JvS, MS and JH conceptualised the comparative genomics analyses. JvS performed the comparative genomics / effectoromics and phylogenetic analyses. JvS and JH wrote the manuscript. JH and GS acquired the main part of the funding and supervised the project. PT, MS, AG, VP and GS sub-stantially revised/commented on the manuscript. All author(s) read the final version of the manuscript and approved it.

---

**Table 1.**
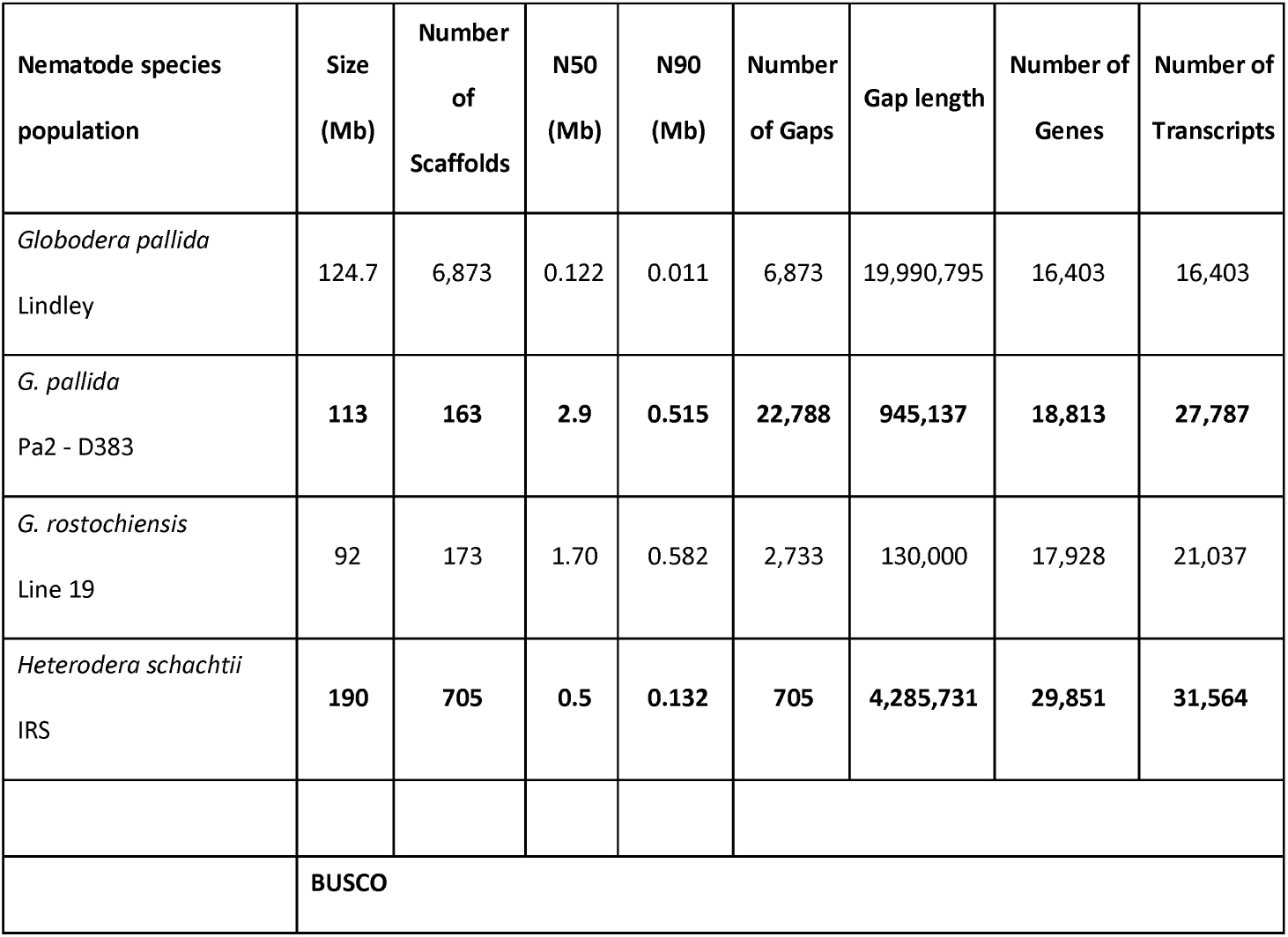
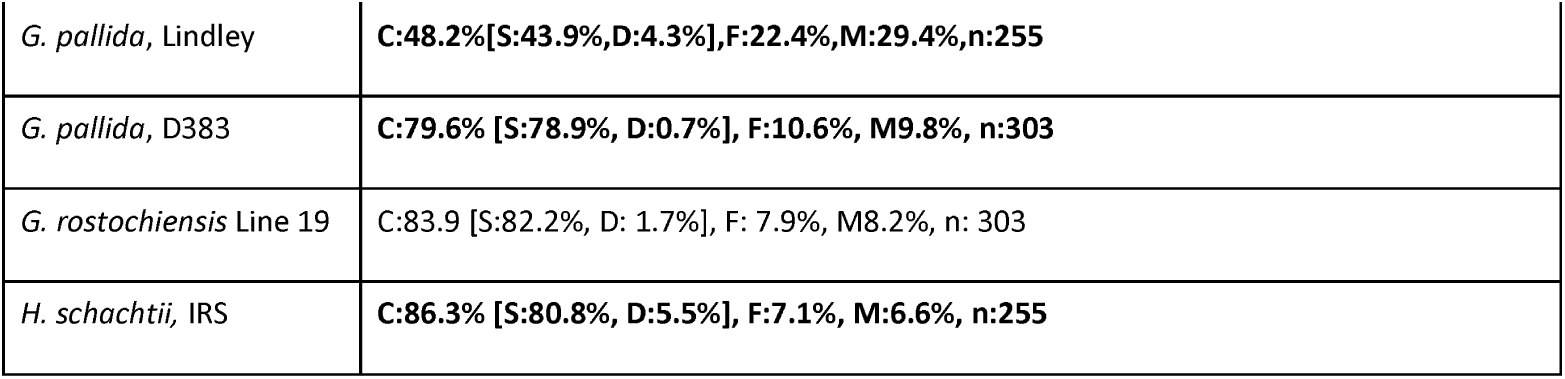
Comparative genome statistics of four cyst nematode genome assemblies. In bold, data from the current paper, data on *Globodera pallida* Lindley and *G. rostochiensis* Line 19 genomes were published by respectively (Cotton et al., 2014) and (Van Steenbrugge et al., 2021)

